# Analysis of SARS-CoV-2 Antibodies in COVID-19 Convalescent Blood using a Coronavirus Antigen Microarray

**DOI:** 10.1101/2020.04.15.043364

**Authors:** Rafael R. de Assis, Aarti Jain, Rie Nakajima, Algis Jasinskas, Jiin Felgner, Joshua M. Obiero, Oluwasanmi Adenaiye, Sheldon Tai, Filbert Hong, Philip J. Norris, Mars Stone, Graham Simmons, Anil Bagri, Martin Schreiber, Andreas Buser, Andreas Holbro, Manuel Battegay, Philip Hosimer, Charles Noesen, Donald K. Milton, Prometheus Study Group, D. Huw Davies, Paul Contestable, Laurence M. Corash, Michael P. Busch, Philip L. Felgner, Saahir Khan

## Abstract

The current practice for diagnosis of COVID-19, based on SARS-CoV-2 PCR testing of pharyngeal or respiratory specimens in a symptomatic patient at high epidemiologic risk, likely underestimates the true prevalence of infection. Serologic methods can more accurately estimate the disease burden by detecting infections missed by the limited testing performed to date. Here, we describe the validation of a coronavirus antigen microarray containing immunologically significant antigens from SARS-CoV-2, in addition to SARS-CoV, MERS-CoV, common human coronavirus strains, and other common respiratory viruses. A comparison of antibody profiles detected on the array from control sera collected prior to the SARS-CoV-2 pandemic versus convalescent blood specimens from virologically confirmed COVID-19 cases demonstrates near complete discrimination of these two groups, with improved performance from use of antigen combinations that include both spike protein and nucleoprotein. This array can be used as a diagnostic tool, as an epidemiologic tool to more accurately estimate the disease burden of COVID-19, and as a research tool to correlate antibody responses with clinical outcomes.

## Introduction

COVID-19 caused by the SARS-CoV-2 virus is a worldwide pandemic with significant morbidity and mortality estimates from 1-4% of confirmed cases^1^. The current case definition for confirmed SARS-CoV-2 infection relies on PCR-positive pharyngeal or respiratory specimens, with testing largely determined by presence of fever or respiratory symptoms in an individual at high epidemiologic risk. However, this case definition likely underestimates true prevalence, as individuals who develop subclinical infection that does not produce fever or respiratory symptoms are unlikely to be tested, and testing by PCR of pharyngeal or respiratory specimens is only around 60-80% sensitive depending on sampling location and technique and the patient’s viral load^2^. Widespread testing within the United States is also severely limited by the lack of available testing kits and testing capacity limitations of available public and private laboratories. Therefore, the true prevalence of SARS-CoV-2 infection is likely much higher than currently reported case numbers would indicate.

Serology can play an important role in defining the true prevalence of COVID-19, particularly for subclinical infection^2^. Early studies of serology demonstrate high sensitivity to detect confirmed SARS-CoV-2 infection, with antibodies to virus detected approximately 1 to 2 weeks after symptom onset^3^. Unlike PCR positivity, SARS-CoV-2 antibodies are detectable throughout the disease course and persist indefinitely^4^. Multiple serologic tests have been developed for COVID-19^5^ including a recently FDA-approved lateral flow assay. However, these tests are limited to detection of antibodies against one or two antigens, and cross-reactivity with antibodies to other human coronaviruses that are present in all adults^6^ is currently unknown. Prior use of serology for detection of emerging coronaviruses focused on antibodies against the spike (S) protein, particularly the S1 domain, and the nucleocapsid protein (NP)^7^. However, the optimal set of antigens to detect strain-specific coronavirus antibodies remains unknown.

Protein microarray technology can be used to detect antibodies of multiple isotypes against hundreds of antigens in a high throughput manner^8,9^ so is well suited to serologic surveillance studies. This technology, which has previously been applied to other emerging coronaviruses^10^, is based on detection of binding antibodies, which are well-correlated with neutralizing antibodies^11^ but do not require viral culture in biosafety level 3 facilities. Recently, our group developed a coronavirus antigen microarray (CoVAM) that includes antigens from SARS-CoV-2 and tested it on human sera collected prior to the pandemic to demonstrate low cross-reactivity with antibodies from human coronaviruses that cause the common cold, particularly for the S1 domain^2^. Here, we further validate this methodology using convalescent blood specimens from COVID-19 cases confirmed by positive SARS-CoV-2 PCR.

## Methodology

### Specimen Collection

A total of 22 de-identified SARS-CoV-2 convalescent blood specimens were collected from nasopharyngeal PCR-positive individuals from different sources with associated data on symptom onset, positive PCR test, and collection (Supplementary Table 1). Two sera were obtained as de-identified discarded laboratory specimens from acute COVID-19 patients from the Oregon Health Sciences University Hospital (OHSU), Portland, OR. These were sourced from discarded clinical laboratory specimens exempted from informed consent and IRB approval under condition of patient anonymity. An additional two sera were obtained from recovered COVID patients at Vitalant Research Institute in San Francisco, CA under an IRB approved protocol. One convalescent plasma was obtained by Cerus Corporation after isolation from a large-volume apheresis collection following standard protocol from a documented recovered COVID-19 blood donor who was more than 28 days post symptomatic. Four plasma samples were obtained from outpatients of the University Hospital Basel, University of Basel, Basel, Switzerland. These patients were screened in accordance with Swiss regulations on blood donation and approved as plasma donors according to the Blood Transfusion Service of the Swiss Red Cross with informed consent. These donors were diagnosed with COVID-19 based on SARS-CoV-2 positive nasopharyngeal swab PCR tests. At time of plasma donation, each had two negative nasopharyngeal swab SARS-CoV-2 PCR tests and negative SARS-CoV-2 PCR tests in blood, and they were qualified as plasma donors. Plasma was collected from these convalescent donors at the Regional Blood Transfusion Service of the Swiss Red Cross in accordance with national regulations.

A total of 144 de-identified pre-pandemic control sera used in this study were collected between November 2018 and May 2019 for a larger study where residents of a college resident community in the Eastern United States were monitored prospectively to identify acute respiratory infection (ARI) cases using questionnaires and RT-qPCR, so as to characterize contagious phenotypes including social connections, built environment, and immunologic phenotypes^12^. Electronic informed consents including future research use authorization was obtained under protocols approved by the Institutional Review Boards (IRBs) of the University of Maryland and the Department of Navy Human Research Protections Office.

### Specimen Testing on Coronavirus Antigen Microarray

The coronavirus antigen microarray used in this investigation includes 67 antigens across subtypes expressed in either baculovirus or HEK-293 cells (Supplementary Table 2). These antigens were provided by Sino Biological U.S. Inc. (Wayne, PA) as either catalog products or custom synthesis service products. The antigens were printed onto microarrays, probed with human sera, and analyzed as previously described^9,13,14^.

Briefly, lyophilized antigens were reconstituted with sterile water to a concentration of 0.1 mg/mL bringing protein solution to 1x phosphate-buffered saline (PBS) and printing buffer was added. Antigens were then printed onto ONCYTE AVID nitrocellulose-coated slides (Grace Bio-Labs, Bend, OR) using an OmniGrid 100 microarray printer (GeneMachines). The microarray slides were probed with human sera diluted 1:100 in 1x Protein Array Blocking Buffer (GVS Life Sciences, Sanford, ME) overnight at 4°C and washed with T-TBS buffer (20 mM Tris-HCl, 150 mM NaCl, 0.05% Tween-20 in ddH2O adjusted to pH 7.5 and filtered) 3 times for 5 minutes each. A mixture of human IgG and IgA secondary antibodies conjugated to quantum dot fluorophores Q800 and Q585 respectively was applied to each of the microarray pads and incubated for 2 hours at room temperature, and pads were then washed with T-TBS 3 times for 5 minutes each and dried. The slides were imaged using ArrayCam imager (Grace Bio-Labs, Bend, OR) to measure background-subtracted median spot fluorescence. Non-specific binding of secondary antibodies was subtracted using saline control. Mean fluorescence of the 4 replicate spots for each antigen was used for analysis.

### Statistical Analyses

The mean fluorescence intensity (MFI) of each antigen was determined by the average of the median fluorescence signal of four replicate spots. The fluorescence signal for each spot was determined by its signal intensity subtracted by the background fluorescence. Antigens containing a human Fc tag were removed from the analysis, as the secondary antibodies used for quantification are known to bind to human Fc; non-human Fc tag did not interfere with the assay. All statistical analyses were conducted using R version 3.6.3 (R Foundation for Statistical Computing, Vienna, Austria).

MFI was normalized by the quantile normalization method using the *proprocessCore* package (version 1.48.0). As a target for normalization, a vector containing the median MFI for IgG or IgA was constructed. Descriptive statistics were used to summarize the IgA and IgG reactivity measured as MFI. Wilcoxon Rank Sum tests with p < 0.05 corrected for multiple comparisons were used to compare the mean differences between groups. In order to rank the antigens from SARS-CoV-2, SARS-CoV, and MERS-CoV for performance in discriminating the positive and negative groups, the Receiver Operating Characteristic Area Under the Curve (ROC AUC) values for each antigen were calculated by comparing positive and negative specimens using the *pROC* package (version 1.16.2). For this, the samples were randomly partitioned into two groups, at a ratio of 75%/25%, using the *caret* package (version 6.9-86). The group with 75% of the samples was used to create a regression model using the *glm* function form the *stats* package (version 3.6.3). The 25% subset was used to predict the outcome of each sample being classified as negative or positive using the *stat* package and the AUC value calculated. This process was repeated for one thousand times and the final AUC values calculated as the median values of all repetitions.

Next, in order to evaluate the benefit of combining antigens for an increased prediction performance, the top ranked antigens, using a cutoff point of auc = 0.85, were combined into groups of all possible combinations from 2 to 4 antigens using the *combinat* package (version 0.0-8). Again, the auc values for each combination were calculated using the same procedure as for individual antigens and the calculated AUCs are representative of the median AUC from one thousand repetitions.

The optimal sensitivity and specificity for each antigen and combination of antigens was calculated based on the maximum Youden Index. Data visualization was performed using the *ggplot2* package (version 3.3.0) or *pROC* package.

## Results

### Construction of Coronavirus Antigen Microarray

A coronavirus antigen microarray (COVAM) was constructed containing 65 antigens that are causes of acute respiratory infections. The array was used to detect IgG and IgA antibodies present in a collection of blood specimens from recovered COVID-19 patients and pre-pandemic control sera, and the results are shown on the heatmap in Figure 1. The viral antigens printed on this array are from epidemic coronaviruses including SARS-CoV-2, SARS-CoV, and MERS-CoV, common cold coronaviruses (HKU1, OC43, NL63, 229E), and multiple subtypes of influenza, adenovirus, metapneumovirus, parainfluenza, and respiratory syncytial virus as listed in Supplementary Table 2. The SARS-CoV-2 antigens on this array include spike protein (S), the receptor-binding (RBD), S1, and S2 domains the whole protein (S1+S2), and the nucleocapsid protein (NP). There is a similar set of antigens represented on the array from SARS-CoV, MERS-CoV, and the four common cold coronaviruses.

**Figure 1.**
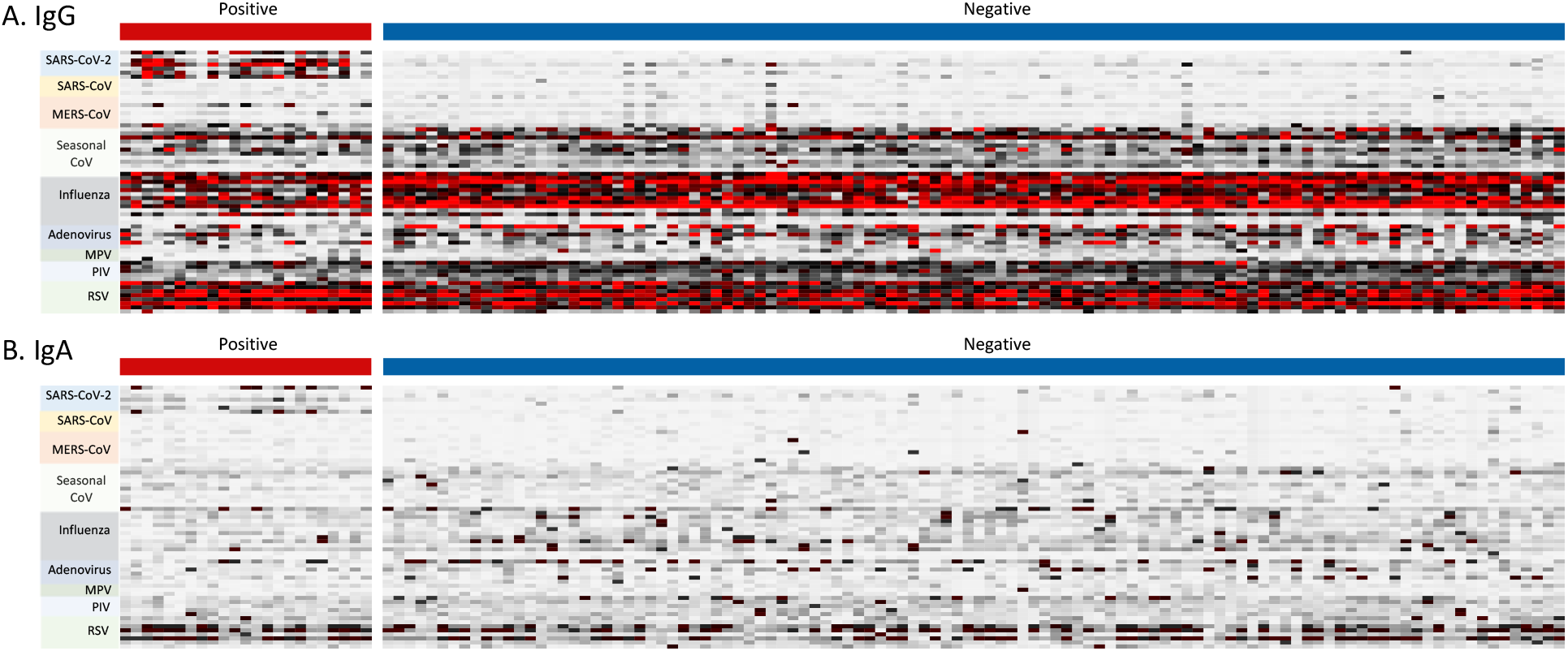
Heatmap for coronavirus antigen microarray. The heatmap shows IgG (A) and IgA (B) reactivity measured as mean fluorescence intensity across four replicates, against each antigen organized into rows color coded by virus, for sera organized into columns classified as positive (convalescent from PCR-positive individuals) or negative (prior to pandemic from naïve individuals). Reactivity is represented by color (white = low, black = mid, red = high).

### Discrimination of SARS-CoV-2 Convalescent Blood Specimens Using Coronavirus Antigen Microarray

To determine the antibody profile of SARS-CoV-2 infection, the differential reactivity to these antigens was evaluated for SARS-CoV-2 convalescent blood specimens from PCR-positive individuals (positive group) and sera collected prior to the COVID-19 pandemic from naïve individuals (negative control group). As shown in the heatmap (Figure 1), the positive group is highly reactive against SARS-CoV-2 antigens. This is more evident for the IgG reactivity then for IgA. The negative controls do not show high reactivity to SARS-CoV-2, SARS-CoV or MERS-CoV antigens despite showing high reactivity to the common cold coronavirus antigens.

With respect to specific antigens, positive group displays high IgG reactivity to SARS-CoV-2 NP, S2, and S1+S2 antigens and to a lesser degree SARS-CoV-2 S1 (Figure 2). The positive group also demonstrates high IgG cross-reactivity against SARS-CoV NP and MERS-CoV S2 and S1+S2 antigens, while the negative group demonstrates low cross-reactivity with S1+S2 and S2 antigens from SARS-CoV-2 and MERS-CoV and no cross-reactivity against other SARS-CoV-2 antigens. The IgA reactivity profile is shown on Figure 3. Overall, IgA seems to follow a similar pattern to IgG, with higher reactivity to SARS-CoV-2 NP, S2 and S1+S2, cross-reactivity to SARS-CoV NP, but no cross-reactivity to the MERS-CoV antigens.

**Figure 2.**
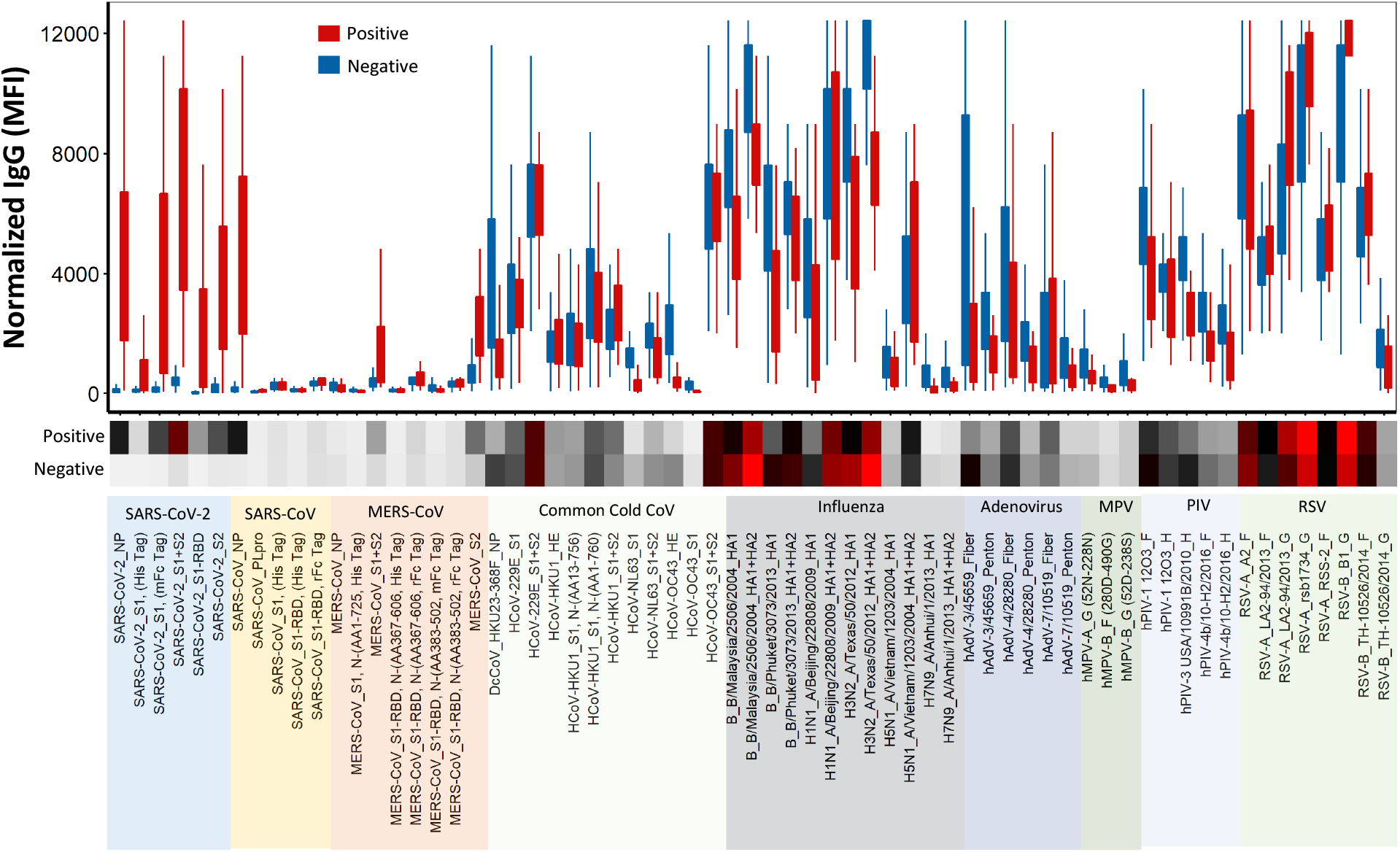
Normalized IgG reactivity of positive and negative sera on coronavirus antigen microarray. The plot shows IgG reactivity against each antigen measured as mean fluorescence intensity (MFI) with full range (bars) and interquartile range (boxes) for convalescent sera from PCR-positive individuals (positive, red) and sera from naïve individuals prior to pandemic (negative, blue). Below the plot, the heatmap shows average reactivity for each group (white = low, black = mid, red = high). The antigen labels are color coded for respiratory virus group.

**Figure 3.**
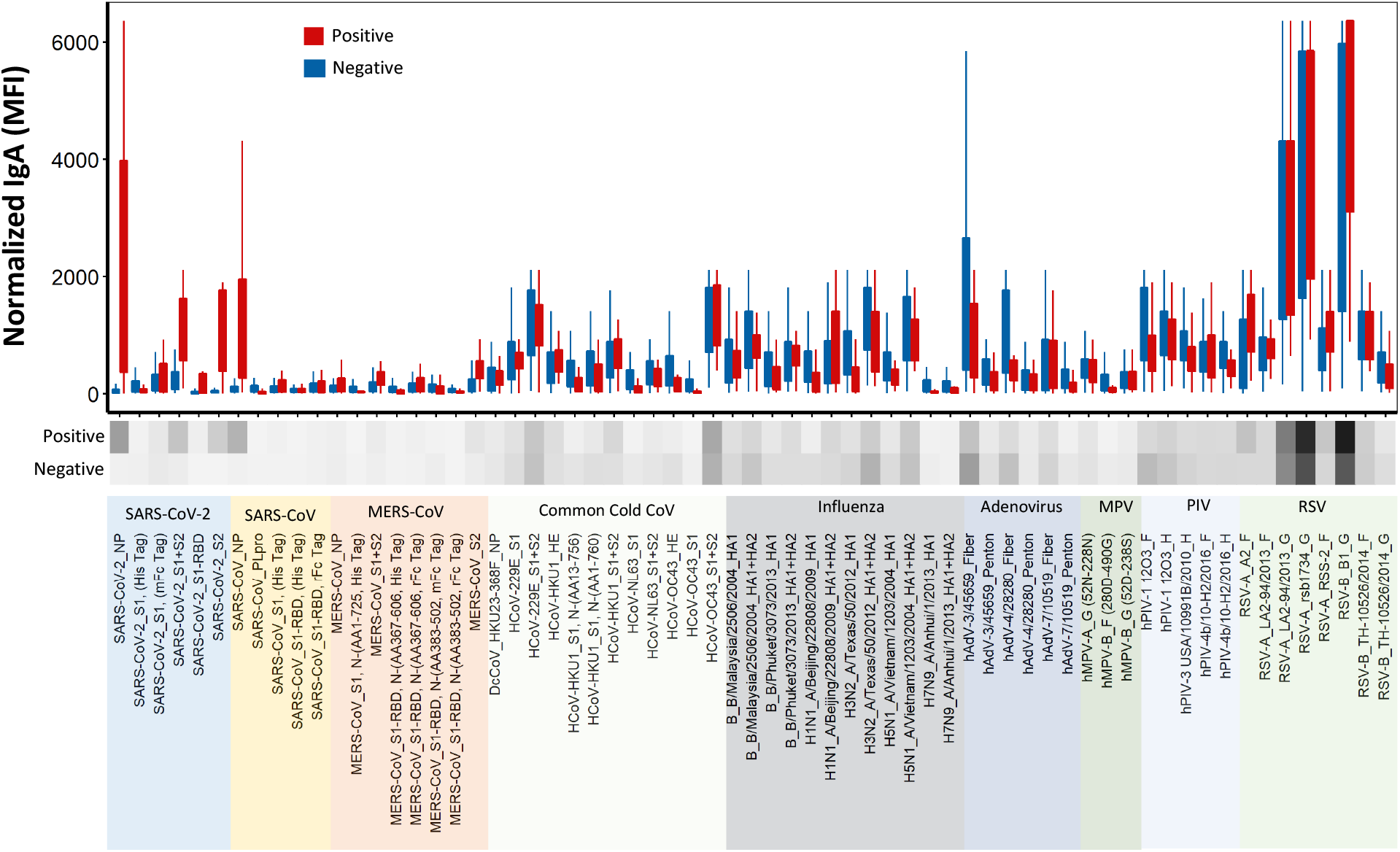
Normalized IgA reactivity of positive and negative sera on coronavirus antigen microarray. The plot shows IgG reactivity against each antigen measured as mean fluorescence intensity (MFI) with full range (bars) and interquartile range (boxes) for convalescent sera from PCR-positive individuals (positive, red) and sera from naïve individuals prior to pandemic (negative, blue). Below the plot, the heatmap shows average reactivity for each group (white = low, black = mid, red = high). The antigen labels are color coded for respiratory virus group.

The two groups do not differ significantly in reactivity to antigens from common cold coronaviruses or other respiratory viruses for either IgG or IgA. The differences between the groups appear to be restricted to SARS-CoV-2 antigens and cross-reactive SARS-CoV and MERS-CoV antigens, so these antigens from epidemic coronaviruses were the focus of subsequent analysis.

### Selection of High-Performing Antigens to Detect SARS-CoV-2 Infection

The sensitivity a specificity of each antigen from SARS-CoV-2, SARS-CoV, and MERS-CoV was evaluated to discriminate the positive group from the negative group across a full range of assay cutoff values using Receiver Operating Characteristic (ROC) curves for which Area Under Curve (AUC) was measured (Figure 4). High-performing antigens for detection of IgG or IgA were defined by ROC AUC ≥ 0.85.

**Figure 4.**
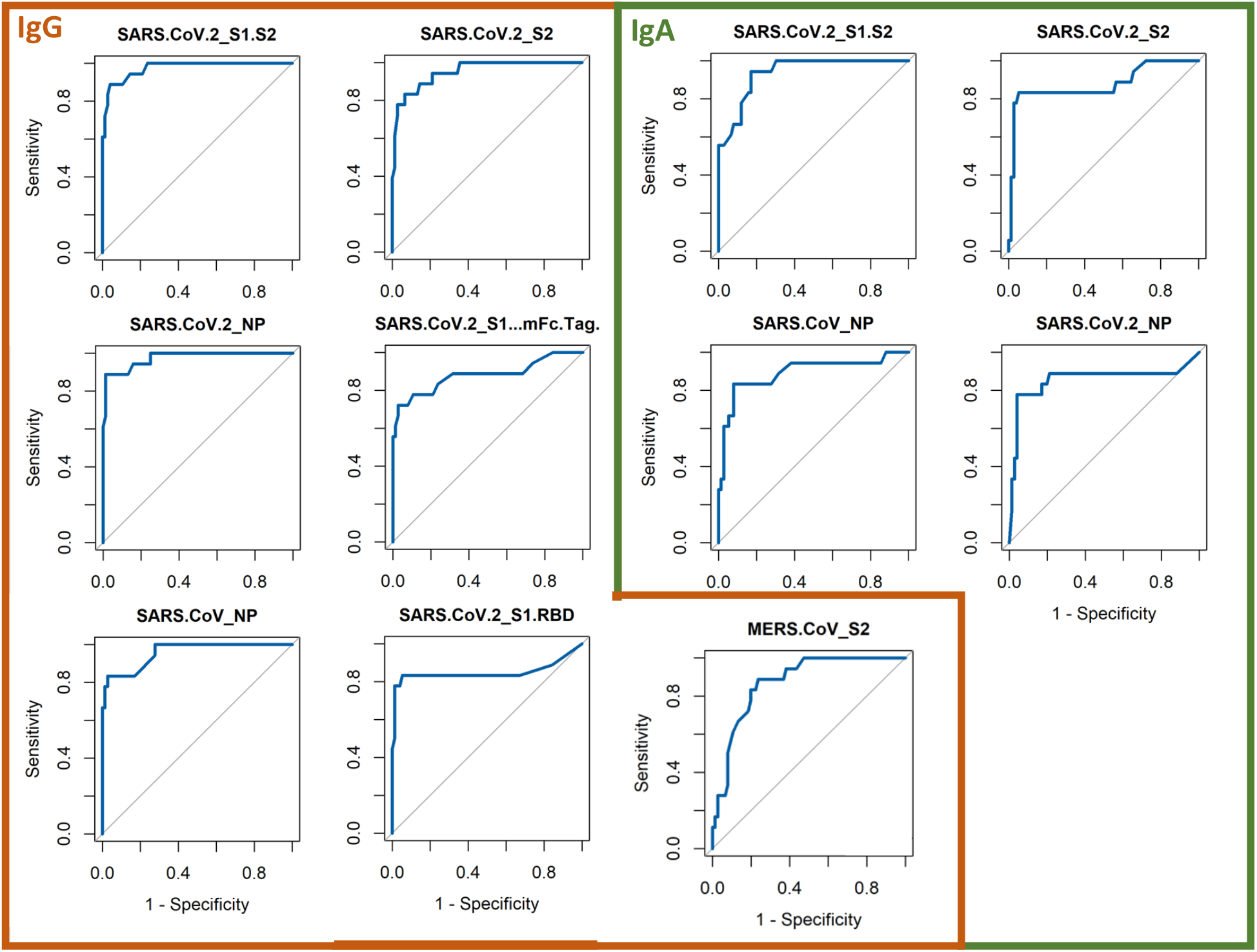
ROC curves for high-performing antigens. ROC curves showing sensitivity versus specificity for discrimination of positive and negative sera were derived for each individual high performing antigen (ROC AUC ≥ 0.95) for both IgG and IgA (solid blue line) and compared to no discrimination (ROC AUC = 0.5, dashed black line).

Although the antigen ranking was different for IgG and IgA, most of the high-ranking antigens were from SARS-CoV-2 and most of the low-ranking antigens were from SARS-CoV and MERS-CoV (Table 1). Among the high-performing antigens, four antigens were ranked as high-performing antigens for both IgG and IgA: SARS-CoV-2 NP, SARS-CoV NP, SARS-CoV-2 S1+S2, and SARS-CoV-2_S2. For IgG, additional high-performing antigens included SARS-CoV-2 S1 (with mouse Fc tag) and RBD and MERS-CoV S2.

**Table 1.** Receiver operating characteristic area under curve (ROC AUC) for SARS-CoV-2, SARS-CoV, and MERS-CoV antigens. ROC AUC values for discrimination of positive and negative sera were derived for each individual antigen for both IgG and IgA and ranked, and high-performing antigens with ROC AUC ≥ 0.86 are indicated above the lines.

| IgG Rank | Antigen | AUC | IgA Rank | Antigen | AUC |
| --- | --- | --- | --- | --- | --- |
| 1 | SARS-CoV-2_S1+S2 | 0.975 | 1 | SARS-CoV-2_S1+S2 | 0.938 |
| 2 | SARS-CoV-2_NP | 0.975 | 2 | SARS-CoV_NP | 0.892 |
| 3 | SARS-CoV-2_S2 | 0.951 | 3 | SARS-CoV-2_S2 | 0.877 |
| 4 | SARS-CoV_NP | 0.957 | 4 | SARS-CoV-2_NP | 0.855 |
| 5 | SARS-CoV-2_S1 (mFcTag) | 0.88 | 5 | MERS-CoV_S2 | 0.803 |
| 6 | MERS-CoV_S2 | 0.873 | 6 | SARS-CoV_PLpro | 0.713 |
| 7 | SARS-CoV-2_S1-RBD | 0.849 | 7 | MERS-CoV_S1.S2 | 0.695 |
| 8 | MERS-CoV_S1+S2 | 0.784 | 8 | MERS-CoV_S1 (AA1-725, His Tag) | 0.677 |
| 9 | SARS-CoV-2_S1 (His Tag) | 0.766 | 9 | SARS-CoV-2_S1 (His Tag) | 0.637 |
| 10 | MERS-CoV_S1-RBD (AA383-502, mFc Tag) | 0.695 | 10 | MERS-CoV_S1-RBD (AA367-606, rFc Tag) | 0.636 |
| 11 | MERS-CoV_S1-RBD (AA367-606, His Tag) | 0.649 | 11 | MERS-CoV_S1-RBD (AA383-502, mFc Tag) | 0.635 |
| 12 | SARS-CoV_PLpro | 0.593 | 12 | SARS-CoV-2_S1-RBD | 0.632 |
| 13 | MERS-CoV_NP | 0.573 | 13 | MERS-CoV_S1-RBD (AA383-502, rFc Tag) | 0.627 |
| 14 | MERS-CoV_S1 (AA1-725, His Tag) | 0.572 | 14 | SARS-CoV-2_S1 (His Tag) | 0.583 |
| 15 | SARS-CoV_S1-RBD (rFc Tag) | 0.510 | 15 | SARS-CoV-2_S1 (mFc Tag) | 0.473 |
| 16 | MERS-CoV_S1-RBD (AA367-606, rFc Tag) | 0.487 | 16 | SARS-CoV_S1-RBD (rFc Tag) | 0.453 |
| 17 | SARS-CoV_S1 (His Tag) | 0.459 | 17 | MERS-CoV_NP | 0.445 |
| 18 | SARS-CoV_S1-RBD (His Tag) | 0.457 | 18 | SARS-CoV_S1-RBD (His Tag) | 0.443 |
| 19 | MERS-CoV_S1-RBD (AA383-502, rFc Tag) | 0.456 | 19 | MERS-CoV_S1-RBD (AA367-606, His Tag) | 0.358 |

Each of the high-performing antigens discriminated between the positive group and the negative group with high significance based on differential reactivity as shown in Figure 5. Positive samples consistently show significantly higher reactivity to these antigens than the negative controls (p < 10^−7^). For IgG, the median reactivity to the top antigens for the positive group is 20-fold higher than for the negative group, while a 6-fold difference is observed for IgA.

**Figure 5.**
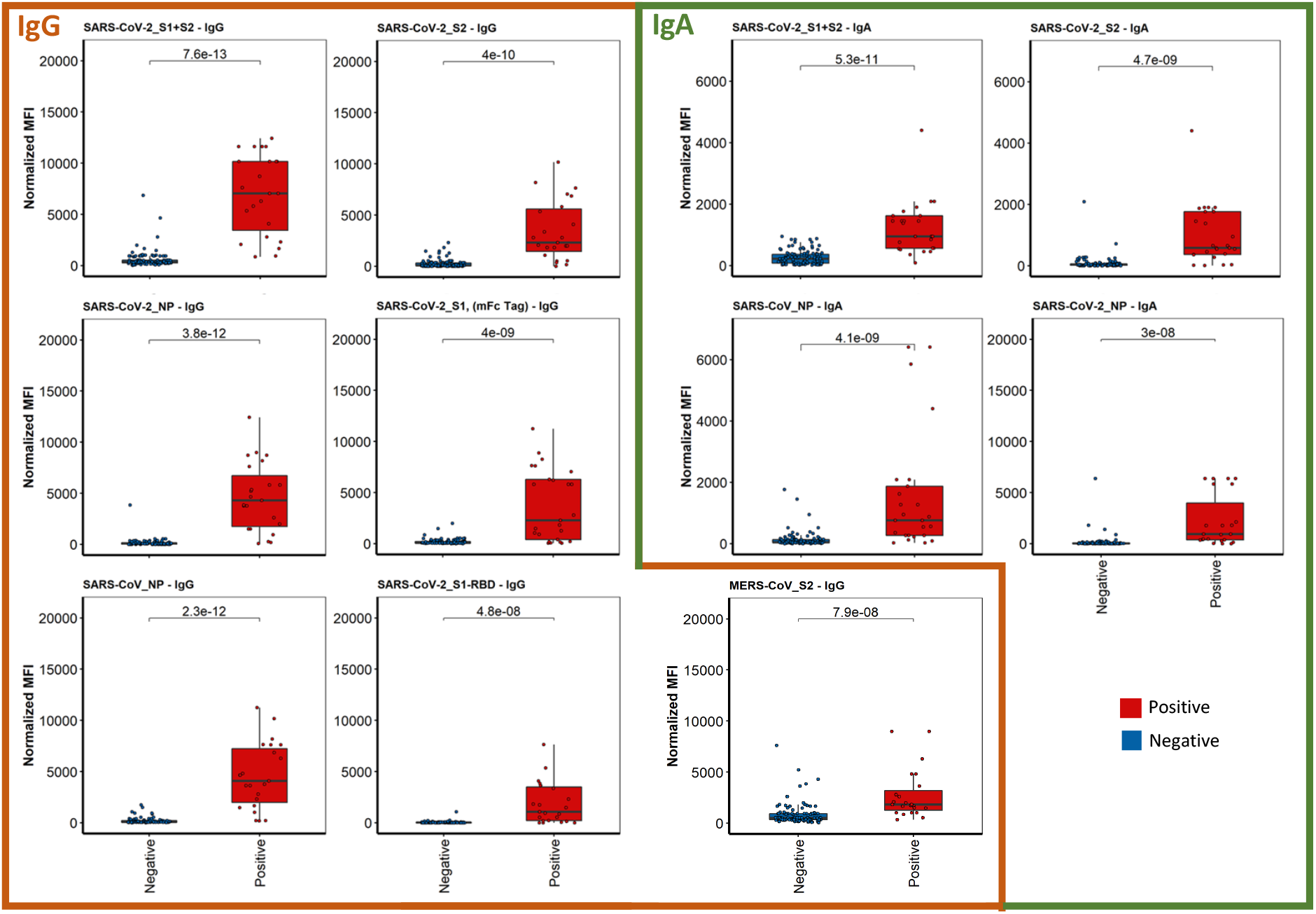
Normalized antibody reactivity of positive and negative sera for high-performing antigens. IgG and IgA reactivity against each high-performing antigens (ROC AUC ≥ 0.95) measured as mean fluorescence intensity (MFI) for convalescent sera from PCR-positive individuals (positive, red) and sera from naïve individuals prior to pandemic (negative, blue) are shown as box plots, including full range (bars), interquartile range (boxes), median (black line), and individual sera (dots) with p-values for each antigen calculated by Wilcoxon Rank Sum test.

The optimal sensitivity and specificity were also estimated for the six high-performing antigens based on the Youden Index (Table 2). For IgG, the lowest sensitivity was seen for SARS-CoV-2 S1, which correlates with the relatively lower reactivity to this antigen in the positive group, while sensitivity was high for the other antigens. The lowest specificity was seen for SARS-CoV-2 S2, which correlates with the cross-reactivity for this antigen seen in a subset of the negative group, while specificity was high for the other antigens. Conversely, for IgA, the highest sensitivity is seen with SARS-CoV-2 S1+S2, and the highest specificity is seen with SARS-CoV-2 S2.

**Table 2.** Performance data for combinations of high-performing antigens. ROC AUC values and sensitivity and specificity based on Youden index for discrimination of positive and negative sera were derived for each individual antigen for both IgG and IgA and ranked, and high-performing antigens with ROC AUC ≥ 0.86 are indicated above the lines.

| N | Antigen Combination | IgG<br>AUC | IgG<br>Spec | IgG<br>Sens | IgA<br>AUC | IgA<br>Spec | IgA<br>Sens |
| --- | --- | --- | --- | --- | --- | --- | --- |
| 1 | SARS-CoV-2_S1+S2 | 0.975 | 0.987 | 0.889 | 0.938 | 0.842 | 0.944 |
| 1 | SARS-CoV-2_NP | 0.975 | 0.961 | 0.889 | 0.855 | 0.947 | 0.778 |
| 1 | SARS-CoV-2_S2 | 0.951 | 0.921 | 0.833 | 0.877 | 0.961 | 0.833 |
| 1 | SARS-CoV_NP | 0.957 | 0.974 | 0.833 | 0.892 | 0.921 | 0.833 |
| 1 | SARS-CoV-2_S1 (mFcTag) | 0.88 | 0.987 | 0.667 | 0.469 | 0.789 | 0.389 |
| 1 | MERS-CoV_S2 | 0.873 | 0.763 | 0.889 | 0.806 | 0.704 | 0.895 |
| 1 | SARS-CoV-2_S1-RBD | 0.849 | 0.947 | 0.833 | 0.645 | 0.974 | 0.444 |
| 2 | SARS-CoV-2_NP ; MERS-CoV_S2 | 0.988 | 0.934 | 1 | 0.914 | 0.934 | 0.833 |
| 2 | SARS-CoV-2_S1+S2 ; SARS-CoV-2_NP | 0.988 | 0.963 | 0.947 | 0.957 | 0.987 | 0.889 |
| 2 | **SARS-CoV-2_S1+S2 ; SARS-CoV_NP** | 0.975 | 0.974 | 0.889 | **0.969** | **0.895** | **0.944** |
| 2 | **SARS-CoV-2_S2 ; SARS-CoV_NP** | **0.994** | **1** | **0.944** | 0.963 | 0.842 | 1 |
| 3 | SARS-CoV-2_NP ; SARS-CoV-2_S2 ; SARS-CoV_NP | 0.988 | 1 | 0.944 | 0.957 | 0.921 | 0.889 |
| 3 | SARS-CoV-2_S1+S2 ; SARS-CoV-2_NP ; SARS-CoV_NP | 0.981 | 1 | 0.889 | 0.963 | 0.816 | 1 |
| 3 | SARS-CoV-2_S1+S2 ; SARS-CoV-2_S2 ; SARS-CoV_NP | 0.975 | 1 | 0.889 | 0.963 | 0.803 | 1 |
| 3 | **SARS-CoV-2_S1+S2 ; SARS-CoV_NP ; SARS-CoV-2_S1, (mFcTag)** | 0.969 | 0.961 | 0.889 | **0.969** | **0.895** | **0.944** |
| 3 | SARS-CoV-2_S2 ; SARS-CoV_NP ; MERS-CoV_S2 | 0.988 | 1 | 0.944 | 0.957 | 0.908 | 0.944 |
| 3 | **SARS-CoV-2_S2 ; SARS-CoV_NP ; SARS-CoV-2_S1, (mFcTag)** | **0.994** | **1** | **0.944** | 0.957 | 0.895 | 0.944 |
| 4 | SARS-CoV-2_S1+S2 ; SARS-CoV-2_NP ; SARS-CoV-2_S2 ; SARS-CoV_NP | 0.981 | 1 | 0.944 | 0.957 | 0.908 | 0.944 |
| 4 | SARS-CoV-2_S1+S2 ; SARS-CoV-2_NP ; SARS-CoV_NP ; SARS-CoV-2_S1-RBD | 0.975 | 1 | 0.833 | 0.963 | 0.816 | 1 |
| 4 | SARS-CoV-2_S1+S2 ; SARS-CoV-2_S2 ; SARS-CoV_NP ; SARS-CoV-2_S1, (mFcTag) | 0.981 | 0.987 | 0.944 | 0.963 | 0.987 | 0.833 |
| 4 | SARS-CoV-2_S1+S2 ; SARS-CoV_NP ; MERS-CoV_S2 ; SARS-CoV-2_S1-RBD | 0.975 | 1 | 0.944 | 0.963 | 0.934 | 0.944 |
| 4 | SARS-CoV-2_S2 ; SARS-CoV_NP ; SARS-CoV-2_S1, (mFcTag) ; SARS-CoV-2_S1-RBD | 0.988 | 1 | 0.944 | 0.944 | 1 | 0.833 |

### Determination of Optimal Antigen Combination to Detect SARS-CoV-2 Infection

In order to estimate the gain in performance by combining antigens, all possible combinations of up to 4 of the 7 high-performing antigens were tested *in silico* for performance in discriminating the positive and negative groups. The ROC curve with AUC, sensitivity, and specificity was calculated for each combination. For both IgG and IgA, there is a clear gain in performance by combining antigens.

The highest performing antigen combinations for each number of antigens are summarized in Table 2, and ROC curves for the top-performing antigen combinations overall for IgG and IgA with comparison to each individual antigen are shown in Figure 6. For IgG, the best discrimination was achieved with the 2-antigen combination of SARS-CoV-2 S2 and SARS-CoV NP, with similar performance upon the addition of SARS-CoV-2 S1 with mouse Fc tag (AUC = 0.994, specificity = 1, sensitivity = 0.944). For IgA, the top performance was achieved with the 2-antigen combination of SARS-CoV-2 S1+S2 and SARS-CoV NP, with similar performance upon the addition of SARS-CoV-2 S1 with mouse Fc tag (AUC = 0.969, specificity = 0.895, sensitivity = 0.944). The addition of a fourth antigen decreased the performance for both IgG and IgA.

**Figure 6.**
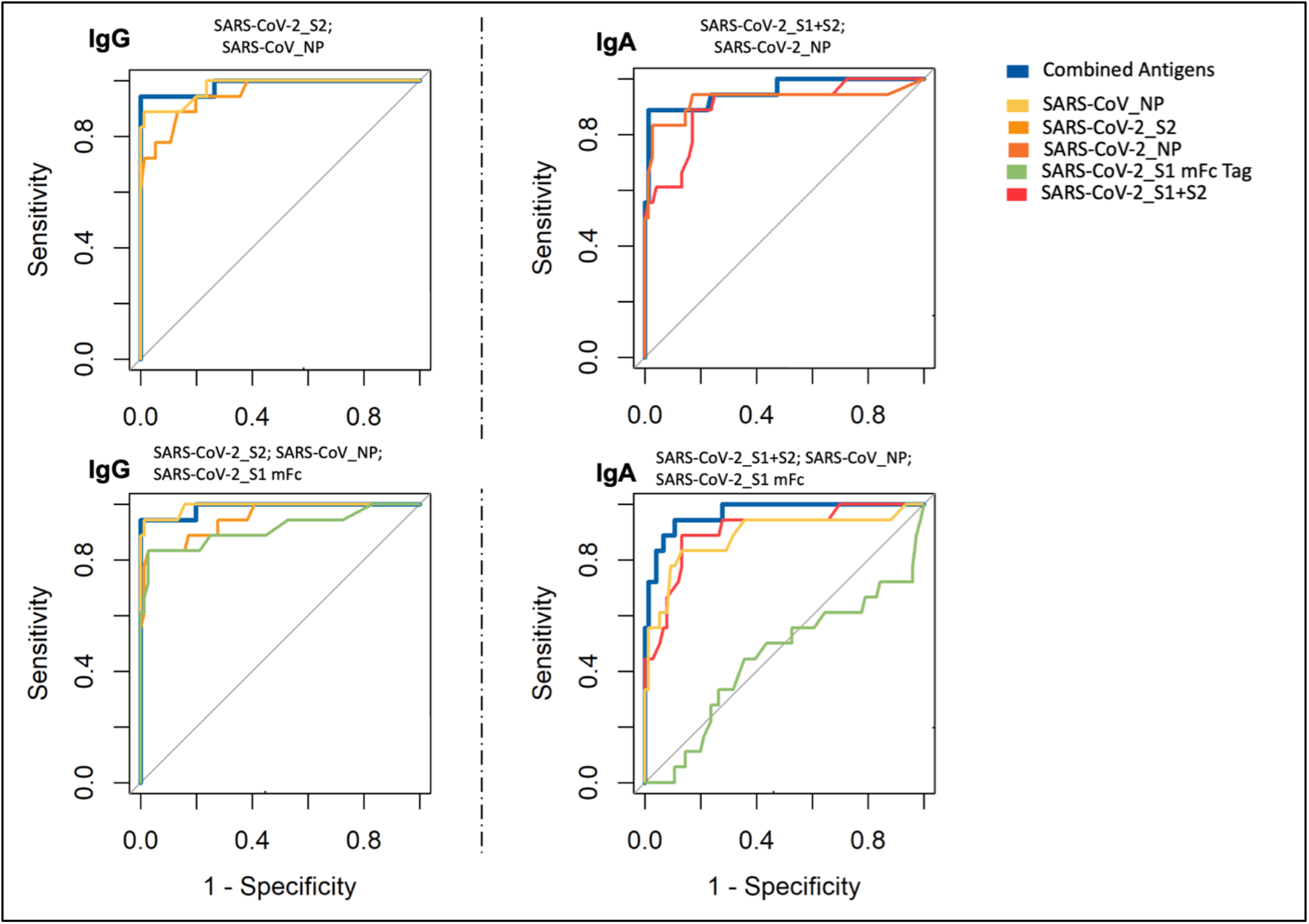
ROC curves for high-performing combination of antigens. ROC curves showing sensitivity versus specificity for discrimination of positive and negative sera were derived for each combination of the high performing antigens for both IgG and IgA (solid blue line) and compared to no discrimination (ROC AUC = 0.5, grey line).

## Discussion

This study reveals several insights into the antibody response to SARS-CoV-2 infection. The antibody profiles of naïve individuals include high IgG reactivity to common cold coronaviruses with low-level cross-reactivity with S2 domains from SARS-CoV-2 and other epidemic coronaviruses, which is not surprising given the high degree of sequence homology and previously observed serologic cross-reactivity^15^ between S2 domains of betacoronaviruses, a group that includes SARS-CoV-2, SARS-CoV, MERS, and common cold coronaviruses HKU1 and OC43. This low-level cross-reactivity occurs in approximately 7% of unexposed individuals (Figure 1), which leads to hypotheses regarding whether these individuals differ in COVID-19 susceptibility and outcomes. However, naïve individuals do not show cross-reactivity to other SARS-CoV-2 antigens. Even for the nucleocapsid protein, which also has high sequence homology between betacoronaviruses, cross-reactivity is only seen between SARS-CoV-2 and SARS-CoV and not with MERS-CoV or common cold coronaviruses. In addition, the quantitative difference between high antibody reactivity to SARS-CoV-2 S2 in the positive group and low-level antibody cross-reactivity in the negative group is large enough that these antigens still discriminate these groups with high significance.

This study also informs antigen selection and design for population surveillance and clinical diagnostic assays and vaccine development. The optimal assay to discriminate SARS-CoV-2 convalescent sera from pre-pandemic sera is a combination of 2 antigens that includes S2 and NP. As an individual antigen, the S2 demonstrates cross-reactivity with negative control sera which leads to low specificity, but this antigen adds predictive power when combined with the more specific NP antigen. The observation that unexposed individuals with antibodies to common cold coronaviruses do not show cross-reactivity to SARS-CoV-2 NP dispels concerns that the high sequence homology of this protein across betacoronaviruses would impair its performance as a diagnostic antigen. The low-level antibody cross-reactivity of a subset of unexposed ndividuals for SARS-CoV-2 spike protein containing S2 domain may not preclude its use as a diagnostic antigen given large quantitative difference in antibody reactivity between positive and negative groups, but this cross-reactivity may influence response to vaccination with spike protein antigens containing the S2 domain in this subset of individuals.

The coronavirus antigen microarray can be useful both as an epidemiologic tool and as a research tool. The high throughput detection of SARS-CoV-2-specific antibody profiles that reliably distinguish COVID-19 cases from negative controls can be applied to large-scale population surveillance studies for a more accurate estimation of the true prevalence of disease than can be achieved with symptom-based PCR testing. In addition, detection of these antibodies in SARS-CoV-2 convalescent plasma donations can provide validation prior to clinical use for passive immunization. The variation in the SARS-CoV-2 antibody profiles among acute and convalescent donors suggests that epitope characterization of convalescent donor plasma will be informative for evaluation of passive immune therapy efficacy in COVID-19 patients. The central role of inflammation in the pathogenesis of severe COVID-19^16^ can be more closely studied by analyzing both strain-specific and cross-reactive antibody responses, particularly to test hypotheses regarding antibody-dependent enhancement with critical implications for vaccine development^17^.

## Conclusions

A coronavirus antigen microarray containing a panel of antigens from SARS-CoV-2 in addition to other human coronaviruses was able to reliably distinguish convalescent plasma of PCR-positive COVID-19 cases from negative control sera collected prior to the pandemic. Antigen combinations including both spike protein and nucleoprotein demonstrated improved performance compared to each individual antigen. Further studies are needed to apply this methodology to large-scale serologic surveillance studies and to correlate specific antibody responses with clinical outcomes.

## Acknowledgements

S. Khan is supported by the National Center for Research Resources and the National Center for Advancing Translational Sciences, National Institutes of Health, through Grant KL2 TR001416. The content is solely the responsibility of the authors and does not necessarily represent the official views of the NIH.

Prometheus-UMD was sponsored by the Defense Advanced Research Projects Agency (DARPA) BTO under the auspices of Col. Matthew Hepburn through agreements N66001-17-2-4023 and N66001-18-2-4015 (PI: D. Milton). This study was funded in part by the Defense Threat Reduction Agency via grants HDTRA1-18-1-0036 (PI: H. Davies) and HDTRA1-18-1-0035 (PI: P. Felgner). The findings and conclusions in this report are those of the authors and do not necessarily represent the official position or policy of the funding agencies and no official endorsements should be inferred.

## Author Contributions

The coronavirus antigen microarray was designed by S. Khan and P. Felgner and was constructed by R. Nakajima and A. Jasinskas at UCI. The negative control sera were collected as research specimens for the Prometheus Study Group by D. Milton, O. Adenaiye, S. Tai, and F. Hong at UMD. The positive COVID-19 convalescent blood specimens were collected as research specimens by L. Corash and A. Bagri at Cerus; M. Busch, G. Simmons, M. Stone, and P. Norris at Vitalant; M. Schreiber at OHSU; P. Hosimer, C. Noesen, and P. Contestable at Ortho Clinical Diagnostics; and A. Buser, A. Holbro, and M. Battegay at University Hospital Basel. The testing of specimens on the coronavirus antigen microarray was performed by A. Jain and J. Felgner at UCI. The data analysis was performed by R. de Assis and J. Obiero at UCI. The manuscript and figures were prepared by S. Khan and R. de Assis with input and approval from all other authors.

## Materials and Correspondence

Please address all correspondence and material requests related to this manuscript to Saahir Khan. The dataset generated by testing specimens on the coronavirus antigen microarray and the analysis code applied to this dataset can be provided upon request.

## Supplementary Information

### Supplementary Tables

**Supplementary Table 1.** SARS-CoV-2 PCR-positive convalescent blood specimens used for validation of the coronavirus antigen microarray. Each de-identified specimen was provided with associated data on symptom onset, positive PCR test, and collection.

| Specimen Source | Specimen Type | Symptom Onset Date | Positive PCR Date | Specimen Collection Date | Days Post Onset |
| --- | --- | --- | --- | --- | --- |
| Vitalant (UCSF) | Serum | 3/11/2020 | 3/13/2020 | 3/30/2020 | 19 |
| Vitalant (UCSF) | Serum | 3/14/2020 | 3/17/2020 | 3/30/2020 | 16 |
| Cerus | Serum | 3/6/2020 |  | 4/4/2020 | 29 |
| OHSU | Serum |  |  |  |  |
| OHSU | Serum |  |  |  |  |
| University Hospital Basel | Plasma | 2/23/2020 | 3/4/2020 | 3/25/2020 | 26 |
| University Hospital Basel | Plasma | 3/3/2020 | 3/8/2020 | 3/27/2020 | 24 |
| University Hospital Basel | Plasma | 3/5/2020 | 3/6/2020 | 3/31/2020 | 26 |
| University Hospital Basel | Plasma | 3/19/2020 | 3/17/2020 | 4/1/2020 | 13 |
| Ortho Clinical Diagnostics | Serum | 3/26/2020 | 3/30/2020 | 4/2/2020 | 7 |
| Ortho Clinical Diagnostics | Serum | 3/20/2020 | 3/26/2020 | 4/2/2020 | 13 |
| Ortho Clinical Diagnostics | Serum | 3/26/2020 | 3/31/2020 | 4/2/2020 | 7 |
| Ortho Clinical Diagnostics | Serum | 3/26/2020 | 3/30/2020 | 4/2/2020 | 7 |
| Ortho Clinical Diagnostics | Serum | 3/20/2020 | 3/27/2020 | 4/2/2020 | 13 |
| Ortho Clinical Diagnostics | Serum | 3/17/2020 | 3/28/2020 | 4/2/2020 | 16 |
| Ortho Clinical Diagnostics | Serum | 3/24/2020 | 3/27/2020 | 4/2/2020 | 9 |
| Ortho Clinical Diagnostics | Serum | 3/14/2020 | 4/1/2020 | 4/2/2020 | 19 |
| Ortho Clinical Diagnostics | Serum | 3/16/2020 | 3/30/2020 | 4/2/2020 | 17 |
| Ortho Clinical Diagnostics | Serum | 3/24/2020 | 3/24/2020 | 4/2/2020 | 9 |
| Ortho Clinical Diagnostics | Serum | 3/24/2020 | 3/28/2020 | 4/2/2020 | 9 |
| Ortho Clinical Diagnostics | Serum | 3/11/2020 | 3/27/2020 | 4/2/2020 | 22 |
| Ortho Clinical Diagnostics | Serum | 3/24/2020 | 3/28/2020 | 3/31/2020 | 7 |
| Ortho Clinical Diagnostics | Serum | 3/24/2020 | 3/27/2020 | 3/31/2020 | 7 |
| Ortho Clinical Diagnostics | Serum | 3/26/2020 | 3/30/2020 | 4/2/2020 | 7 |

**Supplementary Table 2.**
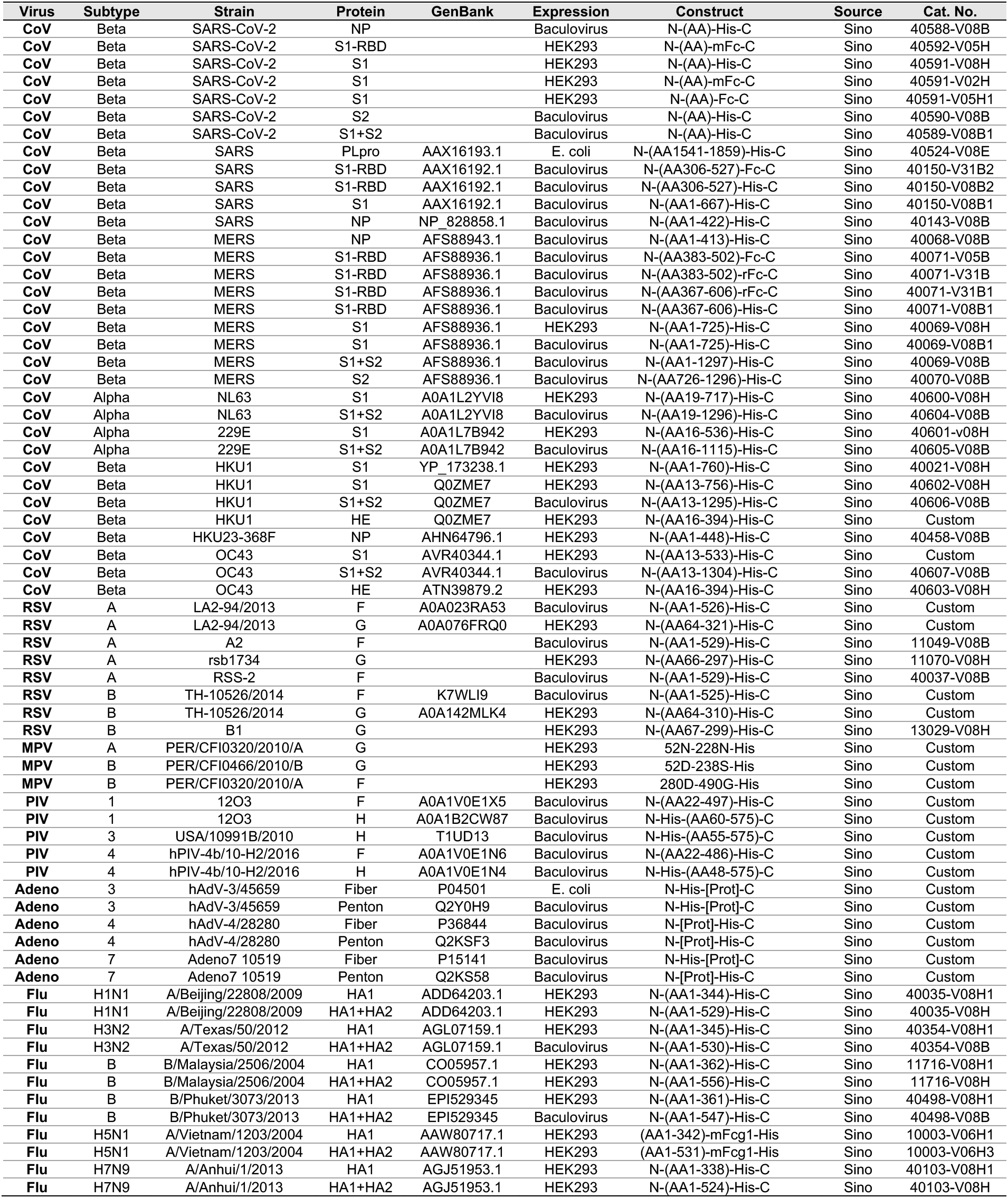
Content of coronavirus antigen microarray. The virus group, subtype, and strain, protein, GenBank identification where available, expression system, gene construct, and vendor source and catalog number are shown for each antigen.

## References

1 Cao, Y., Liu, X., Xiong, L. & Cai, K. Imaging and Clinical Features of Patients With 2019 Novel Coronavirus SARS-CoV-2: A systematic review and meta-analysis. J Med Virol, doi:10.1002/jmv.25822 (2020).

2 Tang, Y. W., Schmitz, J. E., Persing, D. H. & Stratton, C. W. The Laboratory Diagnosis of COVID-19 Infection: Current Issues and Challenges. J Clin Microbiol, doi:10.1128/JCM.00512-20 (2020).

3 Zhao, J. et al. Antibody responses to SARS-CoV-2 in patients of novel coronavirus disease 2019. Clin Infect Dis, doi:10.1093/cid/ciaa344 (2020).

4 To, K. K. et al. Temporal profiles of viral load in posterior oropharyngeal saliva samples and serum antibody responses during infection by SARS-CoV-2: an observational cohort study. Lancet Infect Dis, doi:10.1016/S1473-3099(20)30196-1 (2020).

5 Liu, W. et al. Evaluation of Nucleocapsid and Spike Protein-based ELISAs for detecting antibodies against SARS-CoV-2. J Clin Microbiol, doi:10.1128/JCM.00461-20 (2020).

6 Zhou, W., Wang, W., Wang, H., Lu, R. & Tan, W. First infection by all four non-severe acute respiratory syndrome human coronaviruses takes place during childhood. BMC Infect Dis 13, 433, doi:10.1186/1471-2334-13-433 (2013).

7 Agnihothram, S. et al. Evaluation of serologic and antigenic relationships between middle eastern respiratory syndrome coronavirus and other coronaviruses to develop vaccine platforms for the rapid response to emerging coronaviruses. J Infect Dis 209, 995–1006, doi:10.1093/infdis/jit609 (2014).

8 Davies, D. H. et al. Profiling the humoral immune response to infection by using proteome microarrays: high-throughput vaccine and diagnostic antigen discovery. Proc Natl Acad Sci U S A 102, 547–552, doi:10.1073/pnas.0408782102 (2005).

9 Khan, S. et al. Use of an Influenza Antigen Microarray to Measure the Breadth of Serum Antibodies Across Virus Subtypes. J Vis Exp, doi:10.3791/59973 (2019).

10 Reusken, C. et al. Specific serology for emerging human coronaviruses by protein microarray. Euro Surveill 18, 20441, doi:10.2807/1560-7917.es2013.18.14.20441 (2013).

11 Chan, C. M. et al. Examination of seroprevalence of coronavirus HKU1 infection with S protein-based ELISA and neutralization assay against viral spike pseudotyped virus. J Clin Virol 45, 54–60, doi:10.1016/j.jcv.2009.02.011 (2009).

12 Zhu, S. et al. Ventilation and laboratory confirmed acute respiratory infection (ARI) rates in college residence halls in College Park, Maryland. Environ Int 137, 105537, doi:10.1016/j.envint.2020.105537 (2020).

13 Jain, A. et al. Evaluation of quantum dot immunofluorescence and a digital CMOS imaging system as an alternative to conventional organic fluorescence dyes and laser scanning for quantifying protein microarrays. Proteomics 16, 1271–1279, doi:10.1002/pmic.201500375 (2016).

14 Nakajima, R. et al. Protein Microarray Analysis of the Specificity and Cross-Reactivity of Influenza Virus Hemagglutinin-Specific Antibodies. mSphere 3, doi:10.1128/mSphere.00592-18 (2018).

15 Patrick, D. M. et al. An Outbreak of Human Coronavirus OC43 Infection and Serological Cross-reactivity with SARS Coronavirus. Can J Infect Dis Med Microbiol 17, 330–336, doi:10.1155/2006/152612 (2006).

16 Shi, Y. et al. COVID-19 infection: the perspectives on immune responses. Cell Death Differ, doi:10.1038/s41418-020-0530-3 (2020).

17 Peeples, L. News Feature: Avoiding pitfalls in the pursuit of a COVID-19 vaccine. Proceedings of the National Academy of Sciences, 202005456, doi:10.1073/pnas.2005456117 (2020).

